# Heterogeneous genetic invasions of three insecticide resistance mutations in Indo-Pacific populations of *Aedes aegypti* (L.)

**DOI:** 10.1101/768549

**Authors:** Nancy M. Endersby-Harshman, Thomas L. Schmidt, Jessica Chung, Anthony van Rooyen, Andrew R. Weeks, Ary A. Hoffmann

**Affiliations:** Pest and Environmental Adaptation Research Group, School of BioSciences, Bio21 Institute, 30 Flemington Rd, Parkville, The University of Melbourne, Victoria 3010, Australia; cesar Pty Ltd, 293 Royal Parade, Parkville Victoria 3052, Australia

**Keywords:** insecticide resistance, voltage sensitive sodium channel (*Vssc*), single nucleotide polymorphism (SNP), genetic invasion, linked selection, *Aedes aegypti*

## Abstract

Nations throughout the Indo-Pacific region use pyrethroid insecticides to control *Aedes aegypti*, the mosquito vector of dengue, often without knowledge of pyrethroid resistance status of the pest or origin of resistance. Two mutations (V1016G + F1534C) in the sodium channel gene (*Vssc*) of *Ae. aegypti* modify ion channel function and cause target-site resistance to pyrethroid insecticides, with a third mutation (S989P) having a potential additive effect. Of 27 possible genotypes involving these mutations, some allelic combinations are never seen while others predominate. Here, five allelic combinations common in *Ae. aegypti* from the Indo-Pacific region are described and their geographical distributions investigated using genome-wide SNP markers. We tested the hypothesis that resistance allele combinations evolved *de novo* in populations, versus the alternative that dispersal of *Ae. aegypti* between populations facilitated genetic invasions of allele combinations. We used latent factor mixed-models to detect SNPs throughout the genome that showed structuring in line with resistance allele combinations and compared variation at SNPs within the *Vssc* gene with genome-wide variation. Mixed-models detected an array of SNPs linked to resistance allele combinations, all located within or in close proximity to the *Vssc* gene. Variation at SNPs within the *Vssc* gene was structured by resistance profile, while genome-wide SNPs were structured by population. These results demonstrate that alleles near to resistance mutations have been transferred between populations via linked selection. This indicates that genetic invasions have contributed to the widespread occurrence of *Vssc* allele combinations in *Ae. aegypti* in the Indo-Pacific region, pointing to undocumented mosquito invasions between countries.

## Introduction

Once an invertebrate pest species has invaded a new area, the ability to control the new incursion will depend on whether incursive populations are resistant to chemical pesticides available to control them. This in turn will depend on whether the incursive populations carry pesticide resistance alleles, which can arise through *in situ* evolution of resistance alleles and/or through the introduction of resistance alleles from other established populations. Both processes can be important in pest and disease vector control: examples of local evolution of resistance include pyrethroid resistance in the earth mite *Halotydeus destructor* (Yang et al., 2020) and organophosphate resistance in the whitefly *Bemisia tabaci*, while the long distance introduction of resistance genes is typified by pyrethroid resistance in the mosquito *Culex pipiens* (Chevillon, Raymond, Guillemaud, Lenormand, & Pasteur, 1999); the contribution of both these factors in invading populations is highlighted by pesticide resistance in the spider mite *Tetranychus urticae* (Shi et al., 2019) and the moth *Spodoptera frugiperda* (Nagoshi et al., 2017).

Target-site or knockdown resistance *(kdr)* due to mutations in the sodium channel gene is one of the main mechanisms that compromises control of the dengue vector mosquito, *Aedes aegypti*, with pyrethroid insecticides (Smith, Kasai, & Scott, 2016). These *kdr* mutations have been detected widely in pest insects, following the first discovery of the L1014F mutation in the housefly, *Musca domestica* (Williamson, Denholm, Bell, & Devonshire, 1993), which conferred resistance to DDT. The term *‘kdr’* now covers a range of mutations in different locations in the voltage-sensitive sodium channel (*Vssc)*, with some being found across insect taxa and others being taxon-specific. To aid comparison of mutation sites between taxa, the numbering of codons is usually kept consistent with the codon numbers of the homologous region of the sodium channel gene of the housefly. Although multiple mutations (synonymous and non-synonymous) have been identified in the sodium channel of *Ae. aegypti*, only those non-synonymous mutations found in four positions (codons 1534, 1016, 1011 and 410) have been shown, in electrophysiological assays, to influence the function of the channel so that the toxic action of pyrethroid insecticides is diminished (Du et al., 2013; Haddi et al., 2017) (Figure 1).

**Figure 1.**
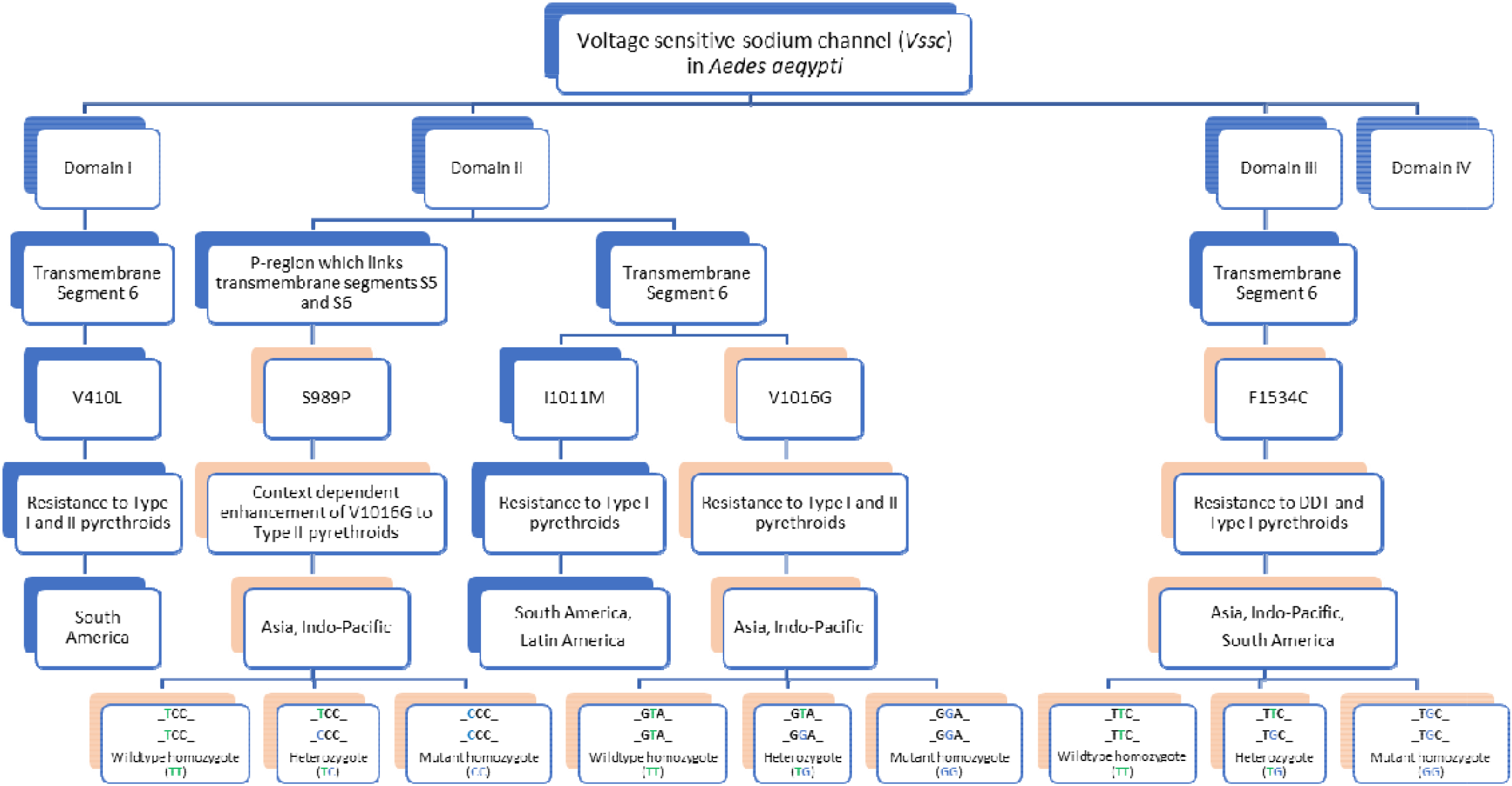
Key to *Vssc* mutations in *Aedes aegypti* that have been functionally characterised and shown to affect the *Vssc* (except for S989P). Codons are numbered according to the homologous sodium channel gene in the house fly, *Musca domestica.* Pink boxes refer to mutations screened in mosquitoes from the Indo-Pacific in this study. Information has been compiled from (Du et al., 2013; Du et al., 2016b; Haddi et al., 2017; Saavedra-Rodríguez et al., 2007; Wuliandari et al., 2015).

Sodium channel mutations at codons 1016 and 1534 have been known for many years in *Ae. aegypti* and occur within the pyrethroid receptor sites in Domains II (S6) and III (S6) of the protein molecule (Du et al., 2013). These two mutations, V1016G and F1534C, are found in *Ae. aegypti* in the Indo-Pacific region (Figure 1). A third mutation, S989P, which is often in perfect linkage with V1016G, is not known to reduce the sensitivity of the sodium channel based on results of Du et al. (2013), but appears to confer some additive pyrethroid resistance in the homozygous state in combination with 1016G in *Ae. aegypti* from Yogyakarta, Indonesia (Wuliandari et al., 2015). S989P is also found in *Ae. aegypti* in the Indo-Pacific region. An additional mutation (D1794Y), which appears to have a similar effect to S989P when found in conjunction with V1016G, is known from *Ae. aegypti tn* Taiwan (Chang, Huang, Chang, Wu, & Dai, 2012; Chang et al., 2009; Lin, Tsen, Tien, & Luo, 2013), and has not been shown to alter the sensitivity of the sodium channel to pyrethroids in electrophysiological assays (Du et al., 2013). A T1520I mutation found in *Ae. aegypti* from India is a third mutation which enhances resistance rather than affecting sodium channel sensitivity by itself and has been shown to increase resistance to Type I pyrethroids caused by F1534C (Chen et al., 2019).

Target-site resistance to pyrethroids is an autosomal, incompletely recessive trait controlled by a single gene (Chang et al., 2012) which has important implications for the resistance status of the heterozygote. Chang et al. (2012) expected the heterozygote at each site to show a level of tolerance to pyrethroid insecticides which is not much higher than that of wildtype (susceptible) individuals and this is the case for S989P+V1016G or F1534C in crossing experiments (Plernsub, Saingamsook, Yanola, Lumjuan, Tippawangkosol, Sukontason, et al., 2016). However, Plernsub et al. (2016) found some enhancement of resistance in the triple heterozygote (S989P/V1016G/F1534C) which showed resistance intermediate between a 1534C homozygous mutant strain and a 989P/1016G homozygous mutant strain in Thailand. Ishak et al. (2015) demonstrated a similar effect in the absence of S989P in *Ae. aegypti* in Malaysia and concluded that V1016G and F1534C heterozygotes occurring in the same individual have an additive effect on deltamethrin resistance of the mosquito.

Crosses performed by Plernsub et al. (2016) between a 1016GG/989PP and a 1534CC strain revealed that combinations of alleles are co-inherited. A haplotype donated by one parent maintains strong linkage patterns between the combination of the mutation sites. This linkage limits genotypes found in offspring of crosses and in the population in general. Several studies have noted that mutant homozygote 1016G is often found in conjunction with a wildtype homozygote at F1534 and *vice versa* (Ishak et al., 2015; Kawada et al., 2014; Stenhouse et al., 2013). Synthesis of data from these studies and our own observations suggest that certain haplotypes of the three mutation sites predominate in a population and there is little evidence of crossing over to disrupt the phase patterns found.

Management of *Ae. aegypti* as a vector of dengue and other arboviruses requires knowledge of its insecticide resistance status and the likelihood of this status changing over time (Moyes et al., 2017; Plernsub, Saingamsook, Yanola, Lumjuan, Tippawangkosol, Walton, et al., 2016). Mosquito populations may become resistant to insecticides by local selection on new mutations or after the incursion of resistant genotypes into a new region which we refer to here as a “genetic invasion”. Local selection pressures from insecticides are expected to vary due to different frequencies of application, rates and proportions of Type I and II pyrethroids applied, and in some cases, will select for specific genotypes, as was observed in *Ae. aegypti* from Yucatan State, Mexico (Saavedra-Rodriguez et al., 2015).

In the absence of strong local selection, resistance alleles are unlikely to increase in frequency in a population because their selective advantage will not be realised and they may carry a fitness cost (Brito et al., 2013). *Vssc* resistance alleles have not been detected in *Ae. aegypti* in northern Australia (Endersby-Harshman et al., 2017) suggesting that they are either not present or occur at very low frequency. If the latter case is correct, then the absence of local selection has prevented resistance alleles from increasing to a detectable frequency in this location. Resistance generated through local selection in one location may spread to others as mosquitoes disperse, resulting in the same resistance mutations occurring in unrelated populations.

Broad-scale geographic variation in the incidence of *Vssc* mutations occurs in *Ae. aegypti;* for example, in Southeast Asia the V1016G mutation is abundant, and in South America the V1016I mutation occurs at the same site; V1016G is clearly causative of pyrethroid resistance while V1016I alone has no effect on sodium channel sensitivity to pyrethroids, but has been shown to increase resistance to both Type I and II pyrethroids in conjunction with the F1534C mutation (Chen et al., 2019). At a local geographic scale, the occurrence and frequency of the 1016, 1534 and 989 mutations can also vary (Leong et al., 2019; C.-X. Li et al., 2015; Wuliandari et al., 2015; Yanola et al., 2011). However, it is not clear if this variation reflects movement patterns of mosquitoes (genetic invasion) or ongoing mutation and selection. Evidence of resistance mutations spreading as mosquitoes move has been found in Ghana where mutation F1534C appeared, as a first record in Africa, associated with an intron phylogeny from Southeast Asia and South America (Kawada et al., 2016) (intron between exon 20 and 21 in the *Vssc* gene).

To resolve how broad-scale resistance patterns are produced, the distribution of resistance alleles can be compared to patterns of genetic differentiation in high resolution single nucleotide polymorphism (SNP) markers (Yang et al., 2020). As the recently developed genome assembly AaegL5 (Matthews et al., 2018) provides a precise genomic location for almost every SNP in *Ae. aegypti*, this methodology can be refined to compare patterns of genetic differentiation in SNPs close to the *Vssc* gene with those of more distant SNPs. Genomic approaches using SNP markers have been effective at identifying differentiation in *Ae. aegypti* across a range of scales (J. E. Brown et al., 2014; Gloria-Soria et al., 2018; Jasper, Schmidt, Ahmad, Sinkins, & Hoffmann, 2019; Rašić, Filipović, Weeks, & Hoffmann, 2014; Schmidt, Filipović, Hoffmann, & Rasic, 2018; Schmidt et al., 2019; Sherpa et al., 2017). The clear differentiation among *Ae. aegypti* populations (Rašić et al., 2014) provides a suitable background against which to compare differentiation at and around the *Vssc* gene region.

Here we report on the distribution of resistance alleles in *Ae. aegypti* from the Indo-Pacific region. We focus on those *Vssc* mutations found in *Ae. aegypti* from the Indo-Pacific region that have been shown in electrophysiological assays to reduce the sensitivity of the mosquito’s sodium channel to pyrethroid insecticides (Du et al., 2013) as well as one mutation which has no effect on channel sensitivity, but may enhance resistance in association with one of the other mutations (Wuliandari et al., 2015). Our study aims to (1) determine the geographical distribution of *Vssc* mutations at codons 989, 1016 and 1534 in *Ae. aegypti* from throughout the region; (2) compare genetic structure at sites near the *Vssc* gene with sites far from the *Vssc* gene; and (3) infer the possible processes leading to the *Vssc* distribution patterns found in mosquitoes throughout the Indo-Pacific.

## Methods

### Insect collection

Samples of Ae. *aegypti* were collected from the field from 43 locations covering 11 countries in the Indo-Pacific region from April 2015 to February 2018 (Supplementary Table S1). Mosquitoes were collected as adults or larvae from water containers and, in the case of larvae, were reared to late instar stages or adults before being confirmed as Ae. *aegypti*, preserved and stored in either >70% ethanol or RNAlater^®^ (AMBION, Inc., Austin, Texas, USA).

### DNA extraction

DNA was extracted from adult mosquitoes or late instar larvae using the DNeasy^®^ Blood and Tissue kit (QIAGEN Sciences, Maryland, USA) according to the instructions of the manufacturer. Two final elutions of DNA were made with the first being used for construction of genomic libraries (see below) and the second being used for screening of *Vssc* mutations after being diluted 1:10 with water.

### *Screening of* Vssc *mutations*

395 samples of *Ae. aegypti* from 11 countries were screened for *Vssc* mutations (Supplementary Table S1). The amino acid positions relating to the mutation sites in this study are labelled as S989P, V1016G and F1534C (Figure 1) according to the sequence of the most abundant splice variant of the house fly, *Musca domestica, Vssc* (GenBank accession nos. AAB476O4 and AAB476O5) (Kasai et al., 2014). These mutation sites are equivalent to those in other studies labelled as S996P, V1023G and F1565C based on the *Vssc* homologue in *Ae. aegypti*, the AaNav protein (GenBank accession no. EU399181) (Du et al., 2013). Custom TaqMan^®^ SNP Genotyping Assays (Life Technologies, California, USA) were developed for each of the three target site mutations (Table 1) based on sequence information from Wuliandari et al. (2015).

**Table 1.**
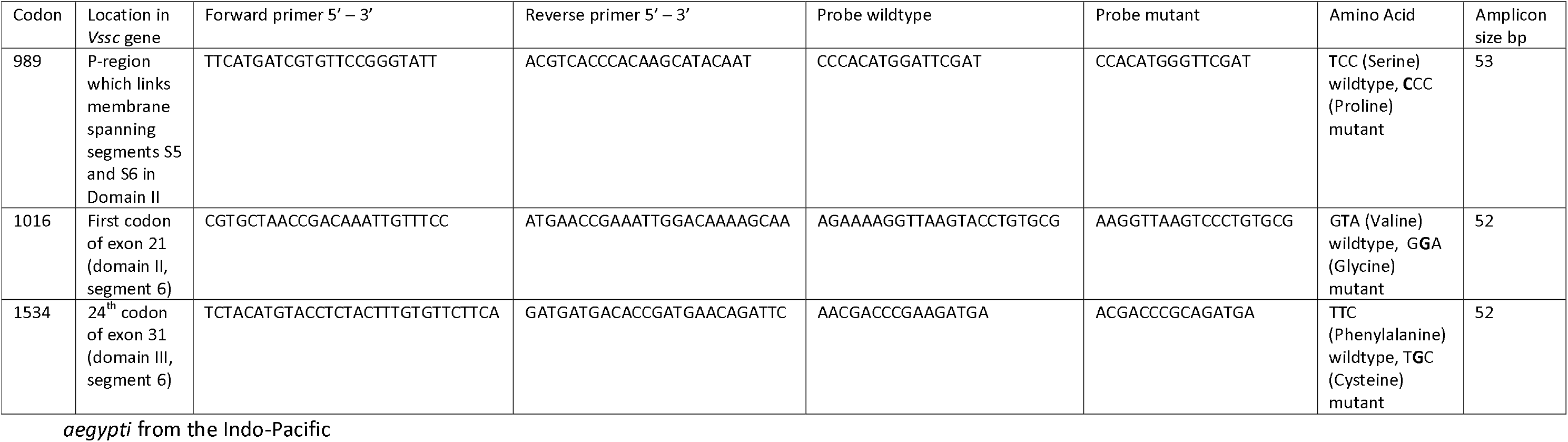
Custom TaqMan^®^ SNP Genotyping Assays (Life Technologies, California, USA) for each of three target site mutations in the *Vssc* gene of *Aedes*

Probes for the wildtype allele in each assay were labelled with Applied Biosystems™ VIC^®^ reporter dye in conjunction with a Minor Groove Binder (MGB) and a non-fluorescent quencher (NFQ). Probes for the mutant allele were labelled with Applied Biosystems™ FAM™ reporter dye, MGB and NFQ. Three replicates of each TaqMan^®^ assay were run on a LightCycler^®^ II 480 (Roche, Basel, Switzerland) real time PCR machine in a 384-well format. The PCR Master Mix contained 4Ox TaqMan^®^ assay as described above (0.174 μL), 2x KAPA Fast PCR Probe Force qPCR Master Mix (KAPABIOSYSTEMS, Cape Town, South Africa) (3.5 μL), ddH_2_O and genomic DNA as prepared above (2 μL). Conditions for the PCR run were pre-incubation of 3 min at 98°C (ramp rate 4.8°C/ s) followed by 40 cycles of amplification at 95°C for 10 s (2.5°C/ s ramp rate) and 60°C for 20 s (2.5°C/ s ramp rate) (Acquisition mode: single) with a final cooling step of 37°C for 1 min (2.5°C/ s ramp rate). Endpoint genotyping was conducted using the Roche LightCycler^®^ 480 Software Version 1.5.1.62.

### Testing for genetic invasions

Tests for genetic invasion were conducted on a single nucleotide polymorphisms (SNP) dataset comprising 80 of the *Ae. aegypti* screened for pyrethroid resistance. We investigated mosquitoes from ten countries, with eight individuals from each country included in the dataset. We considered individuals from the same country to be from the same population. For some populations, more than eight individuals were available for inclusion; in these cases, we first selected individuals for inclusion to preserve maximum variation in *Vssc* genotypes in that population (Figure 2), then selected them in order of having the least missing data. Removing excess individuals ensured that genetic differentiation measurements would not be biased by uneven population sample sizes. For large, genome-wide SNP datasets, five genotypes per population can be sufficient for estimating genetic differentiation (Willing, Dreyer, & van Oosterhout, 2012). We omitted Vietnam from these analyses due to accidental loss of samples, and we also omitted populations lacking resistance mutations. Supplementary Table S2 gives details of the 80 *Ae. aegypti* used to test for genetic invasions. These include some individuals previously sequenced by Schmidt et al. (2019), that have been aligned to the AaegL5 genome assembly (Matthews et al., 2018) used in this study.

**Figure 2.**
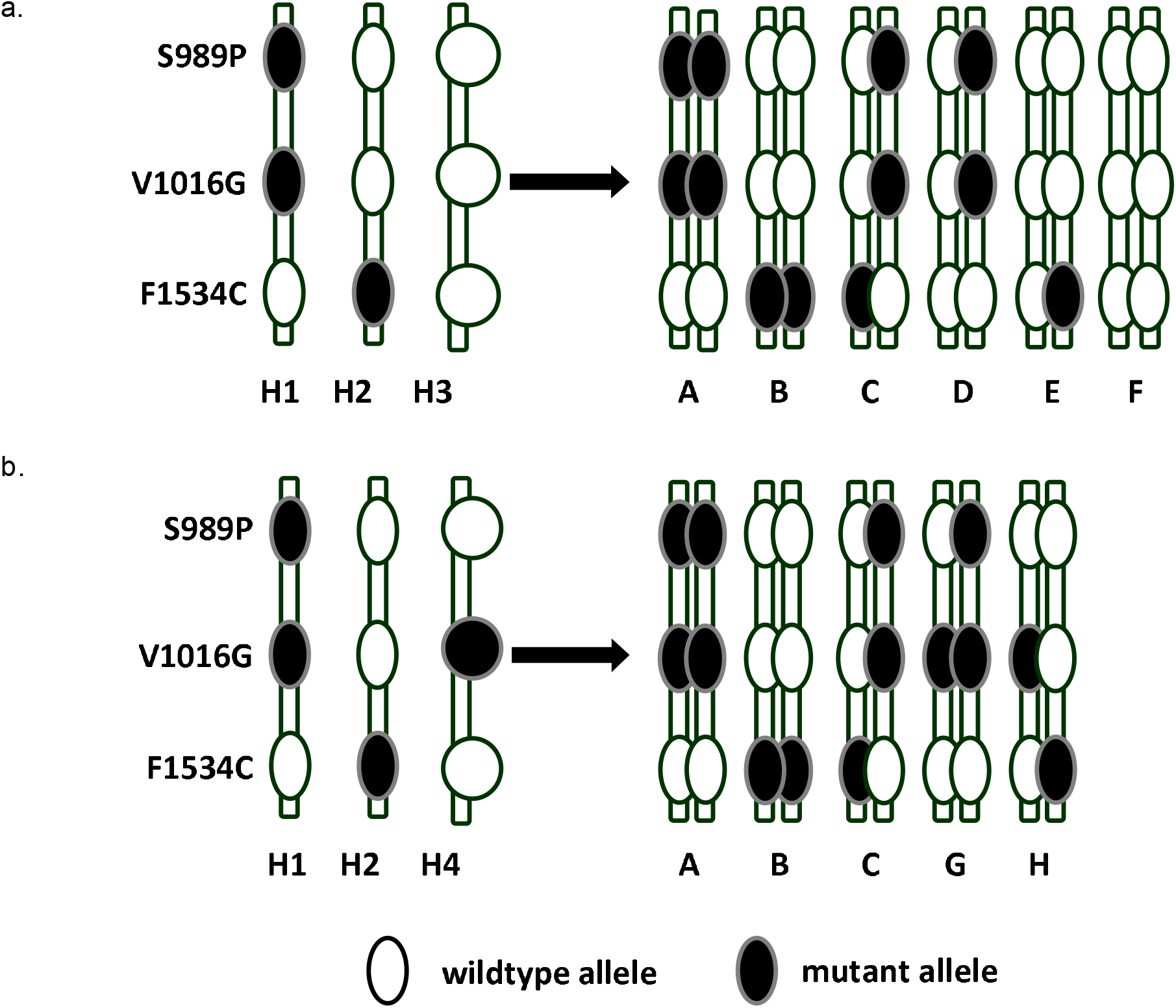
a). Putative haplotypes and phase of common mutant *Vssc kdr* genotypes of *Aedes aegypti* found in the Indo-Pacific region (excluding Taiwan sample), b) Putative haplotypes required to construct genotypes found in Taiwan sample. (Note that H1 = Profile GTC and H2 = Profile TGT).

The 80 *Ae. aegypti* were genotyped for genome-wide SNPs using the double-digest RAD sequencing (ddRADseq) methodology of Rašić et al. (2014) and the bioinformatics pipeline Stacks v2.0 (Catchen, Hohenlohe, Bassham, Amores, & Cresko, 2013). Reads were aligned to the AaegL5 genome assembly (Matthews et al., 2018) using Bowtie2 (Langmead & Salzberg, 2012). The dataset was processed in Stacks and VCFtools (Danecek et al., 2011) with the following filters: SNPs must be biallelic, be present in 75% of mosquitoes in each population, have a minor allele count of ≥ 2 (following Linck and Battey (2019)), have less than 5% missing data, have read depth of between 3 and 45 (following (H. Li, 2014)), and have a known location on one of the three autosomes. The dataset was pre-filtered to ensure no putative first-order relatives were included (all Loiselle’s *k* < 0.1875; (Loiselle, Sork, Nason, & Graham, 1995)). All library construction and filtering steps are detailed in the Supplementary Information S1.

We assigned mosquitoes with the A, C, D, and G genotypes to a resistance profile group “Profile GTC” (one or two copies of haplotype H1; Figure 2), assigned mosquitoes with the B, C, E, and H genotypes to a second group “Profile TGT” (one or two copies of haplotype H2; Figure 2), assigned mosquitoes with the D, E, and F genotypes to a fourth group “Profile TTT” (one or two copies of haplotype H3; Figure 2), and assigned mosquitoes with the G and H genotypes to a third group “Profile GTT” (one or two copies of haplotype H4; Figure 2). Note the nomenclature refers to the mutations in order of degree of resistance conferred from highest to lowest, i.e. 1016/1534/989. Mosquitoes with the C, G, and H genotypes each had two of the above profiles.

To look for indications that resistance alleles have spread via genetic invasion, we investigated patterns of differentiation across the genomes of the 80 individuals. These analyses were motivated by the expectation that, if resistance alleles had been spread by genetic invasion, we would see different patterns of differentiation in SNPs around the *Vssc* gene than at other parts of the genome. Among populations in which a given resistance profile has become established via genetic invasion of a single *de novo* mutation, non-wildtype individuals with identical *Vssc* resistance profiles will have attained these profiles from the same invasive ancestors, and thus these individuals will have alleles that are identical by descent via this invasion. This shared identity by descent should be strongest among alleles at SNPs within and proximate to the *Vssc* gene region (chromosome 3; positions 315,926,360 – 316,405,638), and, specifically, to the point mutation conferring resistance for that profile. Considering the low recombination rate of *Ae. aegypti* (Bennett et al., 2005; S. E. Brown, Severson, Smith, & Knudson, 2001), these patterns of linkage may remain even if gene flow ceased many generations ago or involved only a small number of invasive migrants.

Accordingly, if a given resistance profile had reached its present distribution in the Indo-Pacific region through genetic invasion from a single source population, we would expect to observe similar patterns of variation at SNPs within and proximate to the *Vssc* gene region for individuals with that resistance profile, even when they are from different populations. We would expect variation at other SNPs to be structured by population, as typically observed in *Ae. aegypti* (Rašić et al., 2014; Schmidt et al., 2019). In cases where a population has been recently genetically invaded, there may be very little differentiation in SNPs near the *Vssc* gene among individuals with that resistance profile. Overall, we would expect broad genetic similarity at the *Vssc* gene among all individuals sharing a resistance profile spread via genetic invasion, as alleles near the point mutation actively conferring resistance will be identical by descent. Clearly, analyses should not consider variation at the resistance point mutations themselves, which would confuse any comparison of identity by state and identity by descent. As ddRADseq targets only ~1% of the genome, we considered it unlikely that any of the three resistance point mutations would be sequenced, and we checked to ensure that none of them were.

For a population in which resistance evolved independently, we would also expect to see broad similarities in genetic structure near the *Vssc* gene among individuals in that population sharing that resistance profile. However, while these individuals would have the same resistance mutation identical by state with individuals in other populations, the alleles near the *Vssc* gene would have no identity by descent across populations. Thus, if no genetic invasions have taken place, individuals with the same resistance profile would show structuring in SNPs near the *Vssc* gene along the same lines as SNPs elsewhere in the genome; that is, structuring by population of origin and not by resistance profile.

We performed two analyses to test for genetic invasions, both using the R package “LEA” (Frichot & François, 2015). For the first analysis, we used sparse nonnegative matrix factorization (function *snmf)* to investigate genome-wide patterns of genetic structure and determine an optimal number of clusters *(K)* in which to partition the 80 *Ae. aegypti* genotypes. We then used the optimum *K* to condition a latent factor mixed model (function *lfmm*) (Frichot, Schoville, Bouchard, & François, 2013), which scanned the genome for SNPs that were structured according to a set of environmental variables. For these variables, we used the two resistance profiles: GTC and TGT. We ignored Profile TTT for these analyses; although the susceptible, wild-type profile could serve as an interesting “null” against which to compare the other profiles, almost all of the individuals of Profile TTT were heterozygotes with haplotype H2 (Figure 2), thus preventing any unbiased comparison. Also, as Profile GTT was only found in a single population (Taiwan), it was not appropriate for inclusion in these analyses.

We ran *snmf* using 50 repetitions to determine the value of *K* from 1 to 10 with minimum cross-entropy, which was considered the best estimate of the number of ancestral populations. We ran *Ifmm* with 25 repetitions, using 10,000 iterations and 5,000 burn-in for each. The two resistance profiles were each treated as distinct variables and fit separately. In each case, z-scores were recalibrated using the genomic inflation factor (Frichot & François, 2015).

For the second analysis of genetic invasion, we used the function *pca* to perform two principal components analyses (PCAs) on the 80 *Ae. aegypti*. The first PCA used only the SNPs that were located within the *Vssc* gene region. The second PCA used SNPs that were found anywhere outside the *Vssc* region. In the absence of genetic invasion, we would anticipate that both PCAs would partition the genetic variance roughly equivalently, along population lines. In the case of genetic invasion, we would expect the genetic variance of the *Vssc* SNPs to be structured by resistance profile and that of the non-*Vssc* SNPs by population.

## Results

### Vssc *mutations*

Variation was identified in the *Vssc* alleles and genotype combinations of *Ae. aegypti* throughout the Indo-Pacific region. Eight of the possible 27 combinations of genotypes of the three sodium channel mutation sites were identified in the samples of *Ae. aegypti* and five of these were present in high numbers (Table 2). Mutation sites 1016G and 989P were perfectly linked within our sample with respect to genetic state (except in a small number of individuals from Taiwan) and were in negative linkage disequilibrium with site 1534C. If we consider each possible state, namely wildtype homozygote, mutant homozygote and heterozygote, only three putative haplotypes are required to construct each of the observed combinations (excluding those from Taiwan) (Figure 2a). One additional haplotype is required to construct the two extra genotypes from Taiwan (Figure 2b).

**Table 2.** Frequency of *Vssc kdr* genotypes identified in *Ae. aegypti* from specific countries in the Indo-Pacific region

| **Genotype code:** |  |  | **A** | **B** | **C** | **D** | **E** | **F** | **G** | **H** |
| --- | --- | --- | --- | --- | --- | --- | --- | --- | --- | --- |
| **1016/1534/989:** |  |  | **GGTTCC** | **TTGGTT** | **TGTGTC** | **TGTTTC** | **TTTGTT** | **TTTTTT** | **GGTTTC** | **TGTGTT** |
| **Country** | **n** | **Profile:** | **GTC** | **TGT** | **GTC, TGT** | **GTC** | **TGT** | **TTT** | **GTC** | **TGT** |
| Taiwan | 20 |  | 0.05 | 0.60 | 0.10 |  |  |  | 0.15 | 0.10 |
| Sri Lanka | 24 |  |  | 0.17 |  | 0.04 | 0.63 | 0.17 |  |  |
| Indonesia (Bali) | 67 |  | 1.00 |  |  |  |  |  |  |  |
| Singapore | 29 |  | 0.21 | 0.24 | 0.55 |  |  |  |  |  |
| Malaysia | 30 |  |  | 0.40 | 0.60 |  |  |  |  |  |
| Thailand | 40 |  | 0.25 | 0.50 | 0.25 |  |  |  |  |  |
| Vietnam | 97 |  |  | 0.11 |  |  | 0.47 | 0.41 |  |  |
| Vanuatu | 20 |  | 1.00 |  |  |  |  |  |  |  |
| New Caledonia | 24 |  |  | 0.04 |  |  | 0.33 | 0.63 |  |  |
| Fiji | 24 |  |  | 0.71 |  |  | 0.21 | 0.08 |  |  |
| Kiribati | 20 |  | 0.10 | 0.45 | 0.25 | 0.05 | 0.15 |  |  |  |
| **TOTAL** | **395** |  | **0.27** | **0.23** | **0.13** | **0.01** | **0.19** | **0.15** | **0.01** | **0.01** |

Patterns of *Vssc* mutations were varied across the geographic dataset. From one to five mutation combinations (genotypes) were found at each geographic location with the most combinations (five) being identified from the remote Republic of Kiribati and from Taiwan (Table 2). Mosquitoes from Bali, Indonesia and the Republic of Vanuatu showed only one genotype pattern (homozygous mutant for 1016G and 989P, but wildtype for F1534 – designated as genotype A). Mosquitoes from Vietnam, New Caledonia and Fiji only showed mutations at the 1534 site (homozygous and heterozygous, genotypes B and E) and the sample also contained some completely wildtype individuals (genotype F). *Aedes aegypti* from Singapore, Thailand and Malaysia showed three main genotype combinations, namely A, B and C (C = heterozygous at each of the three sites) (Figure 3).

**Figure 3.**
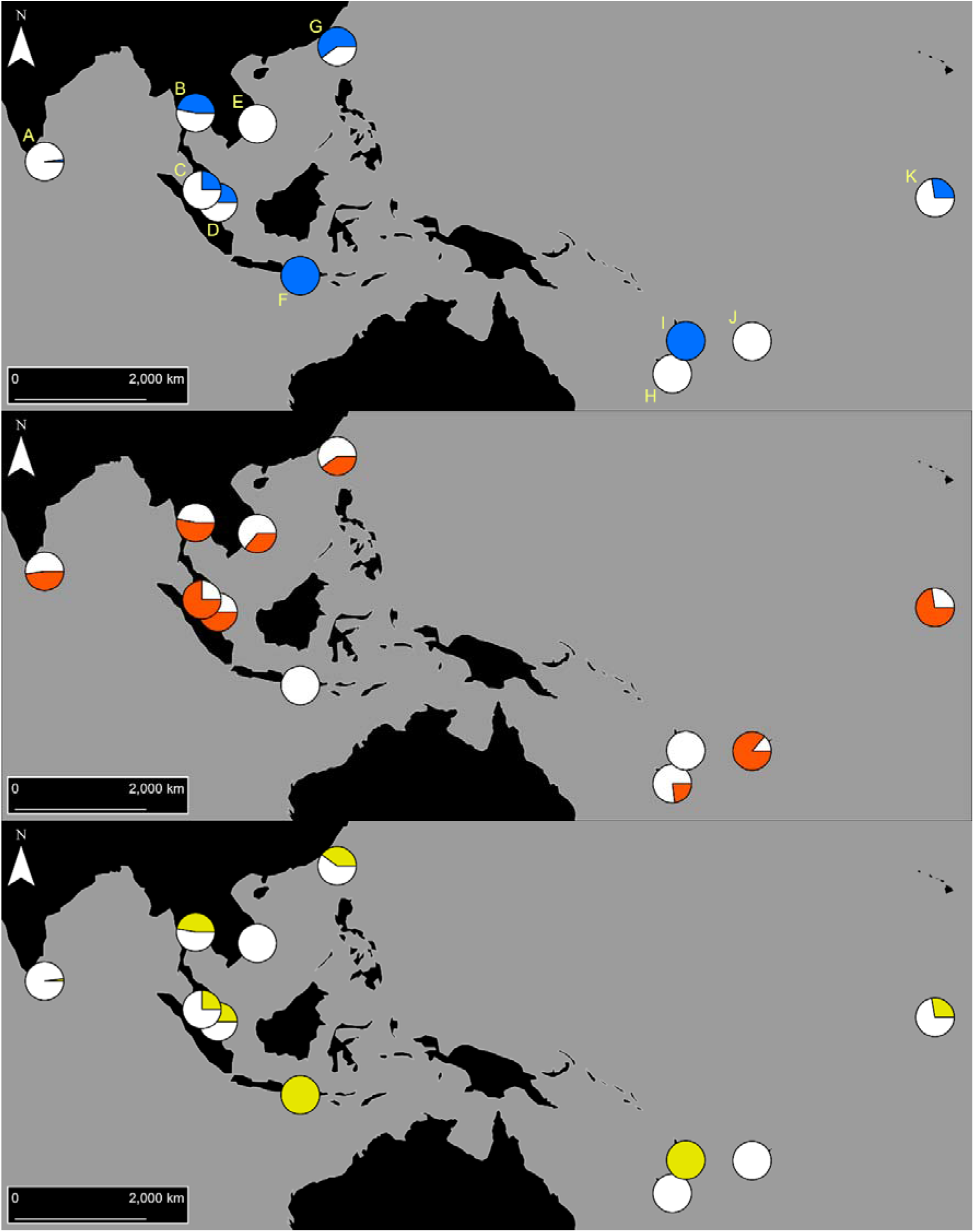
Distribution of the *Vssc* mutations in Indo-Pacific *Aedes aegypti* considered in this study. Coloured sectors indicate the frequency of each type of resistance mutation within each country, following: Blue = 1016G, Orange = 1534C, Yellow = 989P. Populations are: (A) Sri Lanka, (B) Thailand, (C) Malaysia, (D) Singapore, (E) Vietnam, (F) Bali, (G) Taiwan, (H) New Caledonia, (I) Vanuatu, (J) Fiji, and (K) Kiribati.

### Testing for genetic invasions

After filtering, we retained 50,569 genome-wide SNPs for genetic analyses, from an unfiltered set of 93,925 SNPs. Eighteen of these SNPs were located within the *Vssc* gene. Four of these SNPs were located within a *Vssc* exon region, of which three were in the 3’ untranslated region and one was in the coding region. None of the SNPs corresponded to any of the three point mutations conferring resistance. Mean depth across the filtered SNPs was 22.24 (s.d. 6.33) and mean missingness in individuals was 1.82% (s.d. 1.23%).

Using the entire set of 50,569 SNPs, sparse nonnegative matrix factorization in LEA found *K* = 4 to be the optimal choice for *K* (Supplementary Figure S1). Latent factor mixed-models treating Profiles GTC and TGT as environmental factors found a set of SNPs strongly associated with each profile (Figure 4). These SNPs were all clustered around the *Vssc* gene and had -log_10_(P) of up to 57, while elsewhere on the genome no SNP had -log_10_(P) > 15. For Profile GTC, there were 26 SNPs of -log_10_(P) > 15, all of which were found in a region 4,661,744 bp long (chromosome 3; positions 313,105,794 – 317,767,538) surrounding and containing the *Vssc* gene (chromosome 3; positions 315,926,360 – 316,405,638), with eight of these SNPs found within the *Vssc* gene. For Profile TGT, there were 23 SNPs of-log_10_(P) > 15, which were found in a similar region 12,468,872 bp long (chromosome 3; positions 305,090,719–317,559,591) containing the *Vssc* gene. Of these 23 SN Ps, eight were located within the *Vssc* gene, which were the same eight SNPs detected as outliers in the latent factor mixed-model for Profile GTC. Supplementary Figure S2 shows latent factor mixed-model results across a narrow band of chromosome 3 (positions 300,000,000 – 330,000,000).

**Figure 4.**
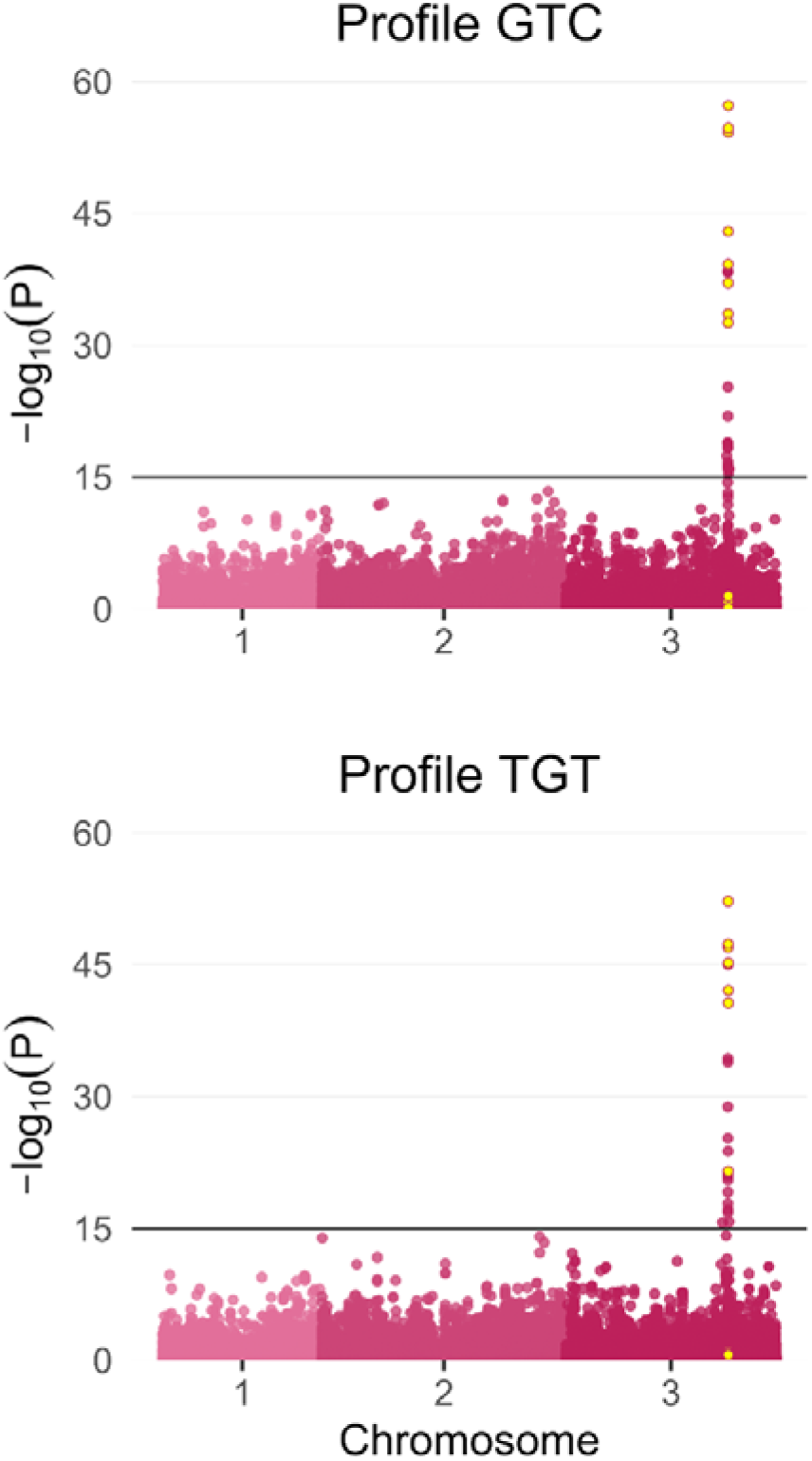
Latent factor mixed-models for Profiles GTC (top) and TGT (bottom). Models use *K =* 4 to condition for population structure among the 80 *Ae. aegypti.* Circles indicate locations and -log_10_(P) of SNPs. Yellow-filled circles indicate SNPs located within the *Vssc* gene (chromosome 3; positions 315,926,360 – 316,405,638).

PCA on the 18 SNPs within the *Vssc* gene region indicated that variation at these SNPs was much more clearly structured by resistance profile than by population (Figure 5a, c, e). When symbology of the PCA was used to indicate individuals of Profile GTC and those not of GTC (Figure 5a), Profile GTC individuals clustered to the top and to the left of most non-GTC individuals. Homozygous GTC individuals (dark blue squares) clustered more tightly and more distinctly than heterozygotes (light blue circles and triangles). A single non-GTC individual from Taiwan (white square) clustered with the 22 GTC homozygotes. Non-GTC individuals with a single copy of the haplotype H4 (green circles) clustered similarly to GTC individuals with a single copy of haplotype H1 (light blue circles and triangles), indicating a potential shared evolutionary origin of these haplotypes. This was also indicated by the single individual with a copy of each of the H1 and H4 haplotypes (light blue circle), which clustered with GTC homozygotes.

**Figure 5.**
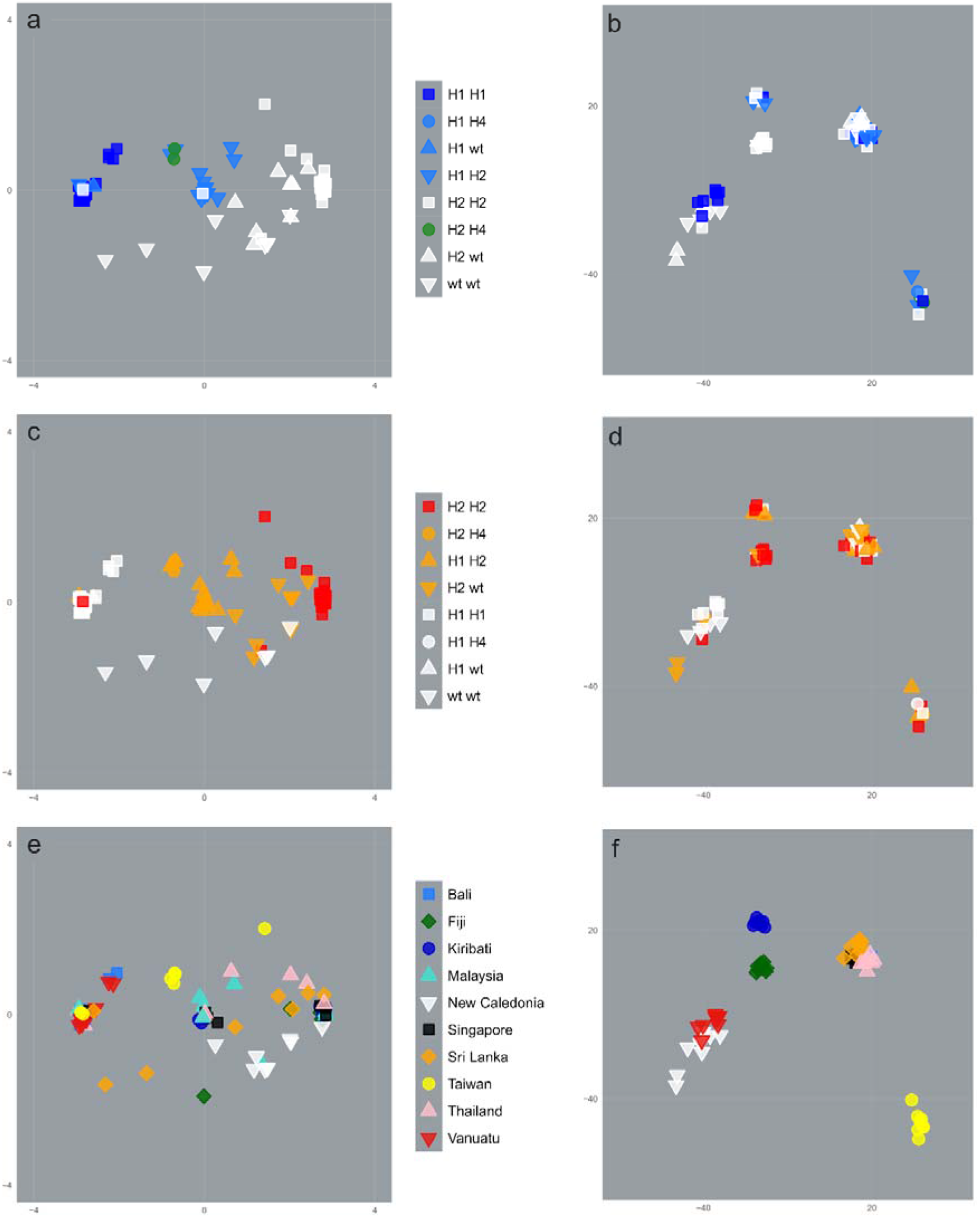
PCAs of 80 *Ae. aegypti* from 10 countries. PCAs show variation at 18 SNPs within the *Vssc* gene (a, c, e) and at 50,551 SNPs outside the *Vssc* gene (b, d, f). PCAs use symbols indicating: individuals of Profile GTC and not of GTC (a, b); individuals of Profile TGT and not of TGT (c, d); and individuals by population (e, f). The green colour used in (a) and (b) reflects the uncertainty surrounding the evolutionary history of haplotype 4. For a, c, and e: PC1 (x-axis) variation = 77.1%, PC2 (y-axis) = 5.9%. For b, d, and f: PC1 (x-axis) variation = 6.7%, PC2 (y-axis) = 4.3%.

When symbology of the PCA was used to indicate individuals of Profile TGT and those not of TGT (Figure 5c), most Profile TGT individuals clustered to the top and to the right of most non-TGT individuals. As with Profile GTC, homozygotes (red squares) clustered more distinctly than heterozygotes (orange circles and triangles). Exceptions were the single Taiwanese homozygote that clustered with GTC homozygotes (white squares), and a Malaysian homozygote that clustered with TGT/wildtype heterozygotes (orange upside-down triangles) and wildtype homozygotes (white upside-down triangles) from New Caledonia.

When symbology of the PCA was used to indicate individuals by population (Figure 5e), no clear structuring was observed among the 18 *Vssc* SNPs. This was most apparent for populations containing a range of resistance profiles, such as Kiribati, Singapore, Sri Lanka, Thailand, and Taiwan. Individuals from these populations were distributed broadly across the PCA plot, indicating a lack of within-population similarity at the *Vssc* gene.

PCA on the 50,551 SNPs outside the *Vssc* gene presented the opposite pattern to that of the *Vssc* gene (Figure 5b, d, f). Among these SNPs, genetic variation was structured unambiguously by population of origin (Figure 5f) and not by resistance profile (Figure 5b, d). The clustering of populations in Figure 5f reflects the *K =* 4 estimated by sparse nonnegative matrix factorization, and shows population separation similar to that observed in previous studies (Schmidt et al., 2019). Clusters were: (1) New Caledonia and Vanuatu; (2) Fiji and Kiribati; (3) Taiwan; and (4) all remaining South and Southeast Asian populations.

Supplementary Figure S3 shows results of additional PCAs. For these, instead of using SNPs found within the *Vssc* gene and those not within the *Vssc* gene, we used SNPs with -logio(P) > 15 associated with Profiles GTC and TGT by latent factor mixed-models (Figure 4). These showed very similar results to Figure 5, wherein variation at -log_10_(P) SNPs was structured by resistance profile (Supplementary Figure S3a, b) and not by population of origin (S2c, d).

## Discussion

This paper has outlined the geographical distributions of three sodium channel mutations found in *Ae. aegypti* from the Indo-Pacific and has presented evidence that these mutations have attained their present regional distributions via genetic invasion. This evidence relates to the genetic structure of SNPs at and around the *Vssc* gene, which showed that mosquitoes from different populations, but with the same resistance profiles, had similar patterns of variation in SNPs near the *Vssc* gene, compared with strong differentiation for the rest of the genome. These patterns indicate that the two widespread *Vssc* genotypes (mutations at codons 1016 and 1534, Profiles GTC and TGT respectively) have spread throughout the region by human transportation of mosquitoes. This addresses an important question about how target-site resistance arises in mosquito populations, which can occur through both genetic invasion and local *de novo* mutation. Our results indicate that strong linkage within the *Vssc* gene region restricts which of the 27 possible genotypes can be produced, which will help inform control strategies suited to local conditions, as resistance status varies greatly among these genotypes. In deriving these results, we also present a detailed geographical summary of pyrethroid resistance in the Indo-Pacific, a critically important region for dengue control.

The negative linkage disequilibrium between *Vssc* mutations at codons 1016 and 1534 as well as the perfect linkage between mutation state at codons 1016 and 989 means that there was no possibility of finding an individual mosquito homozygous for all three mutations. The triple homozygous mutation (989P+1016G+1534C) has been found to enhance resistance to Type I and Type II pyrethroids to a very high level in *Xenopus* oocyte experiments when created artificially in the laboratory (Hirata et al., 2014). Hirata et al. (2014) caution that a single crossing over event could result in an individual with the triple mutation, however, this homozygous combination of mutations has been found only rarely in *Ae. aegypti* from the field (Ishak et al., 2015; Kawada et al., 2014; Sayono et al., 2016) in Penang, Malaysia; Myanmar and Surakarta, Indonesia and its resistance status in that context is not yet clear. Our proposal of inheritance by linked haplotypes with no recombination between mutation sites 1016 and 1534 may explain why the triple mutant does not often arise, even though a heterozygote 1016/1534/989 is common. It is also possible that the triple mutant has a high fitness cost (Hirata et al., 2014; Plernsub, Saingamsook, Yanola, Lumjuan, Tippawangkosol, Sukontason, et al., 2016). A low number of *Ae. aegypti* collected from Saudi Arabia have been identified as being triple mutant heterozygotes with all three mutations on the same chromosome (unlike the combination of our haplotypes H1 and H2, Fig. 2a) and these individuals were susceptible to deltamethrin (Al Nazawi, Aqili, Alzahrani, McCall, & Weetman, 2017).

In the absence of recombination, the presence of only three *Vssc* haplotypes (H1, H2, H3) explains why 21 of the 27 genotypes do not occur in the region (excluding Taiwan). The addition of a fourth haplotype (H4) in the sample from Taiwan enables formation of the two extra genotypes (G and H, Fig 2) found there. If H4 were combined with H3, another of the 27 possible genotypes could be constructed, but individuals of this genotype (TG/TT/TT) were not observed. The new haplotype in Taiwan (H4) may be a result of recombination, with the H1 haplotype found elsewhere in the world. It may be that this haplotype was introduced to Taiwan and then recombined to produce H4. Alternatively, H1 and H4 could both have evolved in Taiwan, before the H1 haplotype spread elsewhere, although it is then not clear why H4 has failed to spread. The effect of the genotypes G and H that we found only in Taiwan on pyrethroid resistance is not known. Genotype G has also been recorded in *Ae. aegypti* in Myanmar (Kawada et al., 2014), but resistance levels of mosquitoes with this genotype were not tested.

Our analyses of resistance Profiles GTC and TGT indicate that genetic invasion is likely to have played a significant role in establishing the current distributions of their associated haplotypes. Our latent factor mixed-models detected regions surrounding the *Vssc* gene 4,661,744 and 12,468,872 bp long in which there were 26 and 23 SNPs closely associated with resistance profiles (Figure 4).

Investigating SNPs within the *Vssc* region using PCA also showed these were structured by resistance profile rather than by population (Figure 5). As alleles at these SNP loci should not be conferring any selective advantage related to resistance, we can conclude that their structuring is a result of linked selection with SNPs conferring resistance. We would expect to see these SNPs structured according to population if resistance mutations had arisen *de novo* in populations. Instead, we see clear evidence that haplotypes H1 and H2 have genetically invaded the Indo-Pacific region. While the role of human transportation in establishing geographical distributions of *Aedes* mosquitoes is already well recognised (Tatem, Hay, & Rogers, 2006), these results indicate that dispersal along transportation networks has also helped to establish current distributions of pyrethroid resistance mutations. These results stand in contrast to a similar investigation of pyrethroid resistance in Australian red-legged earth mite *(Halotydeus destructor)* populations, which showed that the present distribution of resistance had been attained by multiple *de novo* mutations (Yang et al., 2020).

We propose that a series of genetic invasions have established haplotype H1 in Bali, Kiribati, Malaysia, Singapore, Sri Lanka, Thailand, and Vanuatu, and have established haplotype H2 in Fiji, Kiribati, Malaysia, Singapore, Sri Lanka, Thailand, and Vietnam. The consistent structuring by resistance profile of *Vssc* genetic variation from these populations accords with every copy of these haplotypes sharing identity by descent. However, there is no indication from our results whether either haplotype originated in any of these populations or elsewhere within the Indo-Pacific region, which will require further sampling. For instance, we were not able to include samples from Hawaii, Tahiti, or the Philippines, three Indo-Pacific locations with interesting patterns of genetic structure relative to other Indo-Pacific populations (Gloria-Soria et al., 2016).

From our results, it seems likely that Taiwan and New Caledonia have also developed local resistance following the same genetic invasions, though these populations both had some resistant individuals that did not cluster convincingly with others having the same resistance profile (Figure 5). In Taiwan, there were two such cases: one, the clustering of Haplotype H4 along similar lines as H1 (Figure 5a), and another, the clustering of a single Profile TGT homozygote with Profile GTC individuals (Figure 5c). The first observation is best explained by the H1 and H4 haplotypes having a shared evolutionary origin, with one being the ancestral haplotype and one being derived. Determining which is ancestral is beyond the scope of this study and would, at minimum, require more widespread sampling throughout the Indo-Pacific region. The second observation is less easily explained, but potentially relates to one or more recombination events within the *Vssc* gene introducing variation associated with the H1 haplotype into the H2 haplotype in Taiwan. A more comprehensive investigation of resistance in Taiwanese *Ae. aegypti* will be necessary to resolve these issues. The imperfect clustering of New Caledonia and haplotype H2 (Figure 5c) appears to relate more to the rareness of resistance haplotypes in New Caledonia. Only five copies of H2 were observed there, compared with 11 copies of the wild-type H3 haplotype, which likely prevented strong structuring by resistance profile. The solitary resistant homozygote (genotype B, Figure 2) from New Caledonia did cluster convincingly with other resistant homozygotes, presenting evidence for genetic invasion in this population.

Populations of *Ae. aegypti* studied by others have shown evidence of selective sweeps affecting this gene around codons 1534 (Ishak et al., 2015) and 1016 (Wuliandari et al., 2015). Similar evidence of selection has been found in *Ae. aegypti* from South America around codon 1016 (Saavedra-Rodriguez et al., 2007), but segregation of South American mosquitoes from those of the Indo-Pacific region has long been recognized (Smith et al., 2016), due, in particular, to the geographic restriction of V1016I to South America. Saavedra-Rodriguez et al. (2007) showed high levels of recombination between codons 1011 and 1016 in *Ae. aegypti* from South America, but mutations at 1011 have are not been recorded in *Ae. aegypti* from the Indo-Pacific or southeast Asia (Kawada et al., 2009; Kawada et al., 2014; Smith et al., 2016). Evidence for local selection of pyrethroid resistance has been obtained in *Ae. aegypti* from Mexico (Saavedra-Rodriguez et al., 2015). V1016G appears to have arisen before S989P (C.-X. Li et al., 2015), as S989P is never (Kawada et al., 2014) or rarely (Wuliandari et al., 2015) found alone. Our data indicate that some sites have likely reached a stable point of resistance (e.g. Bali, Vanuatu) given that genotypes are fixed. However, others are in flux and we expect that the resistance we have recorded may change in the future. A recent study of *Ae. aegypti* in Taiwan (Biduda et al., 2019) has shown an increase over time in frequency of the 1534C mutation, indicating that this population is still undergoing change.

Smith et al. (2016) summarised the findings of multiple studies on the combinations of mutations that lead to pyrethroid resistance as follows: the V1016G mutation found alone confers resistance to Type I (those without an α-cyano group) and Type II pyrethroids (those with an α-cyano-3-phenoxybenzyl group) whereas F1534C alone confers resistance only to pyrethroids of Type I and to DDT (Du, Nomura, Zhorov, & Dong, 2016a). The degree of resistance induced by V1016G alone is four times that of F1534C alone when expressed in *Xenopus* oocytes (Hirata et al., 2014). S989P alone in *Xenopus* oocyte trials has not been shown to confer resistance (Hirata et al., 2014). The geographic haplotype distribution we have observed in *Ae. aegypti* from the Indo-Pacific suggests that mosquito control in Bali and Vanuatu will be very difficult with both Type I and Type II pyrethroids. Some pyrethroid efficacy is likely to have been lost to various degrees in Taiwan, Kiribati, Malaysia, Singapore, Sri Lanka and Thailand. Mosquito control in Fiji, Vietnam and New Caledonia with Type I pyrethroids may be compromised, but Type II pyrethroids are likely to remain effective.

The potential impacts of resistance conferred by *Vssc* mutations, however, may be modified by other factors. For example, Du et al. (2016b) noted context dependent effects of combinations of *Vssc* mutations likely related to genetic background of the mosquito. Smith et al. (2019) found that a combination of *kdr* and CYP-mediated metabolic detoxification of insecticides confers a greater than additive level of resistance to *Ae. aegypti*, so there may be a selective advantage to mosquitoes having both mechanisms. In that case, the spread of *kdr* mutations that we observed may have been accompanied by the spread of other resistance mechanisms.

Our approach of looking at the *Vssc* mutations in the context of genetic population structure helps indicate the extent of movement of resistance alleles in *Ae. aegypti* in the Indo-Pacific and provides little evidence for independent evolution of pyrethroid resistance in different populations throughout the region. The results of this study likely reflect a series of genetic invasions that have proceeded from the initial biological invasions of the Indo-Pacific by *Ae. aegypti* (J. E. Brown et al., 2014). These invasions have introduced sets of allele combinations conferring resistance to insecticides used in the region. Our results point to the importance of biosecurity controls to prevent resistance alleles moving to new areas with mosquito incursions. Insecticide resistance pressures within a country need to be reduced in order to prevent resistance alleles becoming fixed in mosquito populations.

## Acknowledgements

We thank our mosquito collectors – Ashley Callahan, Jason Axford, Tim Hurst, Elizabeth Valerie, Craig Williams. The study was funded by the Department of Agriculture and Water Resources (DAWR), Australian Government; National Health and Medical Research Council with a Programme Grant/Award Numbers: 1037003, 1132412; a Fellowship to Ary A. Hoffmann, NHMRC Fellowship Grant no. 1118640 and the Wellcome Trust, UK. This research was facilitated by use of the Nectar Research Cloud, a collaborative Australian research platform supported by the Australian Research Data Commons (ARDC) and National Collaborative Research Infrastructure Strategy (NCRIS).

## Data Accessibility Statement

Resistance genotype data are available within the manuscript. The aligned .bam sequence files are available through the Sequence Read Archive at NCBI Genbank, BioProject ID PRJNA608612.

## Author Contributions

NMEH, TLS, AAH designed the study. NMEH, TLS, JC performed research and analysed data. AvR contributed assay design. NMEH, TLS, AAH, JC and ARW wrote the paper.

