## Supplemental Figure S2 for "Heterogeneous genetic invasions of three insecticide resistance mutations in Indo-Pacific populations of *Aedes aegypti* (L.)"

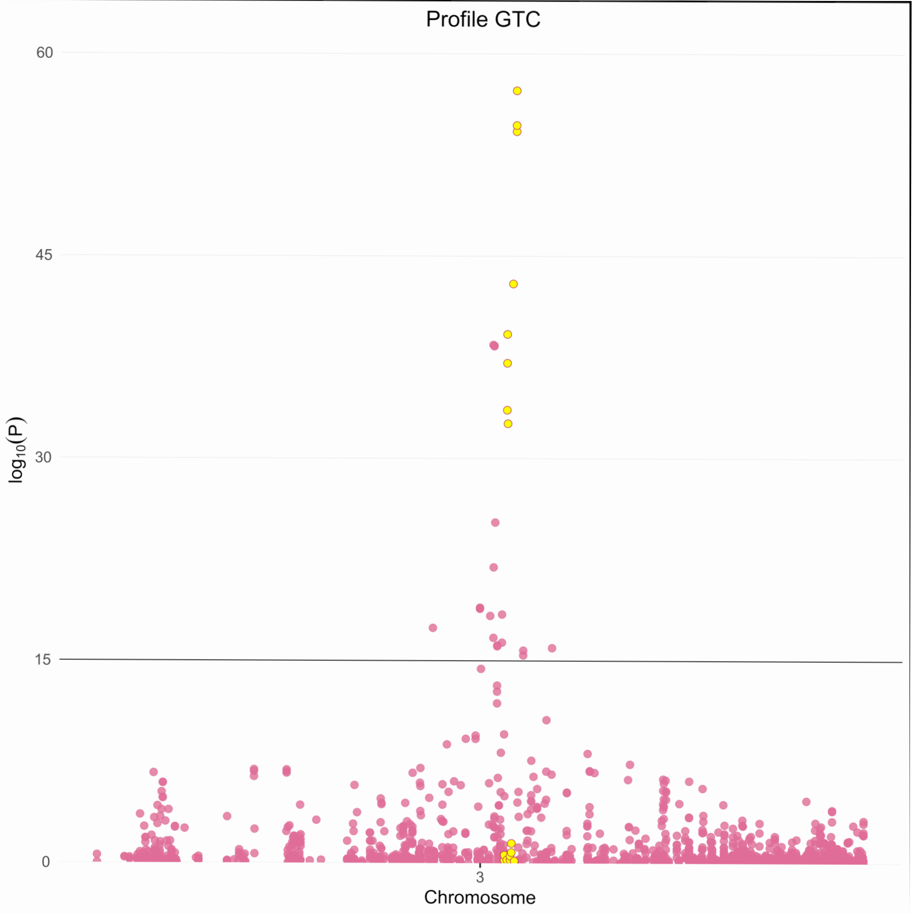
Profile GTC
60
45
30
15
0
log10(P)
3
Chromosome

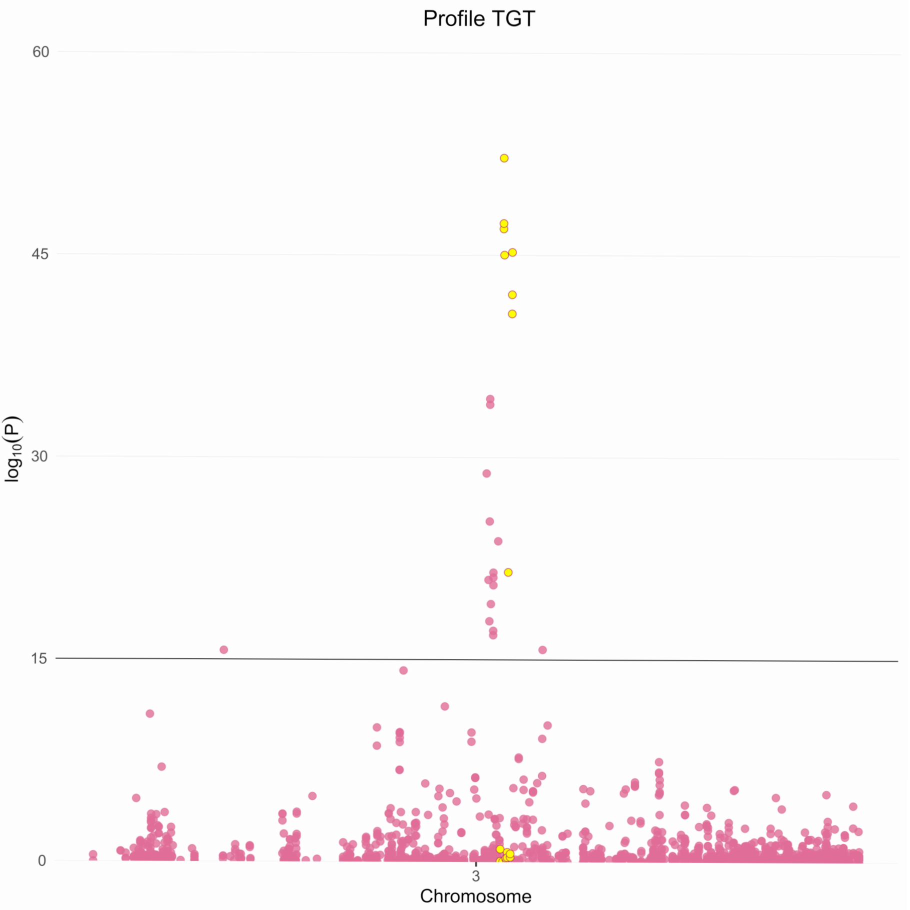
Profile TGT
60
45
30
15
0
log10(P)
3
Chromosome

Supplementary Figure S2. Latent factor mixed-models results across a narrow band of chromosome 3

(positions 300,000,000 – 330,000,000).

From:
Heterogeneous genetic invasions of three insecticide resistance mutations in Indo-Pacific populations of *Aedes aegypti* (L.)

Nancy M. Endersby-Harshman[1*●], Thomas L. Schmidt[1*], Jessica Chung[1], Anthony van Rooyen[2], Andrew R. Weeks[1,2], Ary A. Hoffmann[1]

[1]Pest and Environmental Adaptation Research Group, School of BioSciences, Bio21 Institute, 30 Flemington Rd, Parkville, The University of Melbourne, Victoria 3010, Australia
[2]cesar Pty Ltd, 293 Royal Parade, Parkville Victoria 3052, Australia

*Equal first authors
