## Supplemental Figure S3 for "Heterogeneous genetic invasions of three insecticide resistance mutations in Indo-Pacific populations of *Aedes aegypti* (L.)"

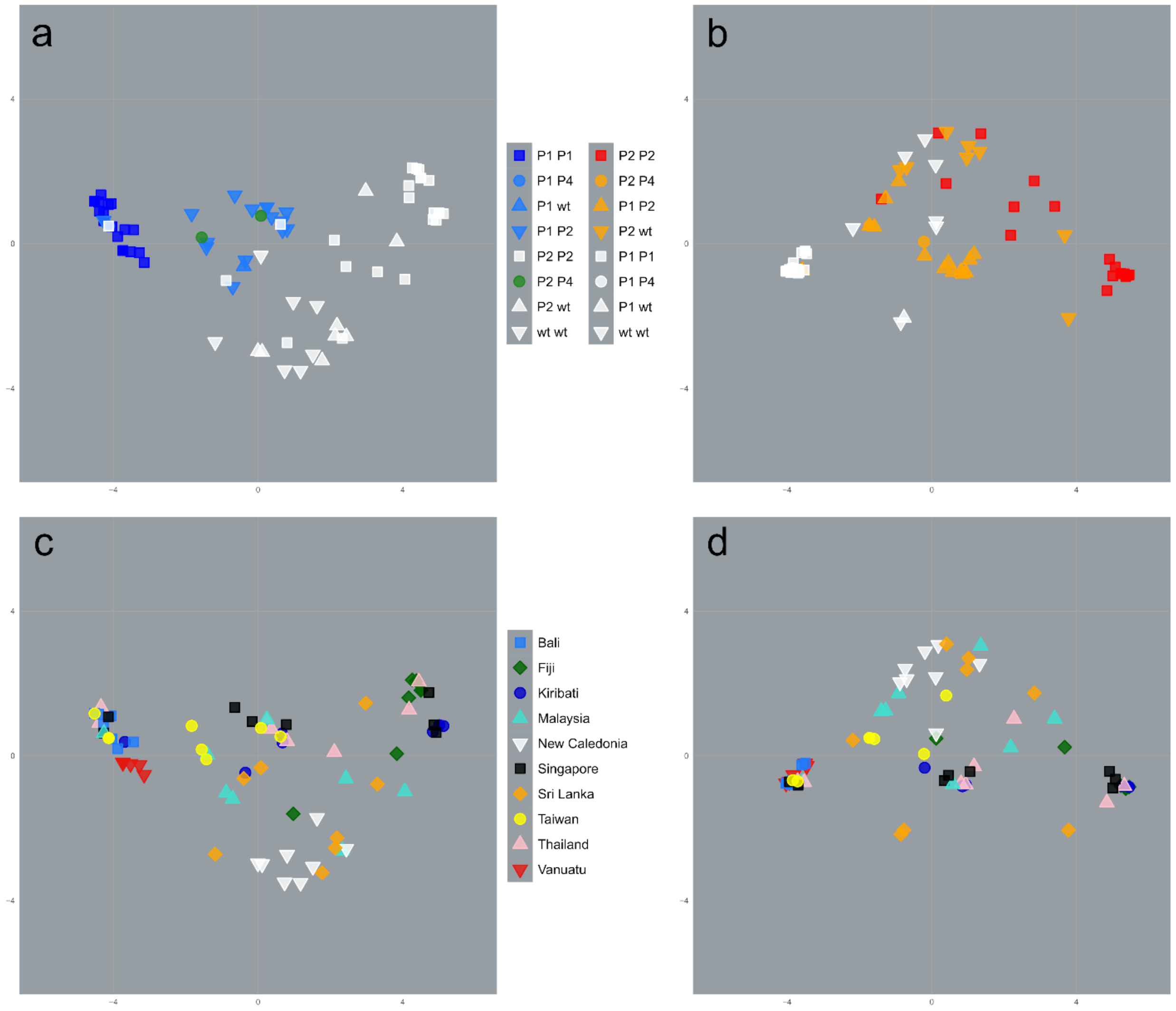


**Supplementary Figure S3. PCAs of 80 *Ae. aegypti* from 10 countries.**

PCAs show variation at 26 SNPs with $-\log_{10}(P) > 15$ associated with Profile GTC by latent factor mixed-model (a, c) and at 23 SNPs with $-\log_{10}(P) > 15$ associated with Profile TGT by latent factor mixed-model (b, d). PCAs use symbolisation indicating: individuals of Profile GTC and not of GTC (a); individuals of Profile TGT and not of TGT (b); and individuals by population (c, d). The green colour used in (a) reflects the uncertainty surrounding the evolutionary history of haplotype 4. For a and c: PC1 (x-axis) variation = 58.1%, PC2 (y-axis) = 12.0%. For b and d: PC1 (x-axis) variation = 68.9%, PC2 (y-axis) = 10.4%.

From:

**Heterogeneous genetic invasions of three insecticide resistance mutations in Indo-Pacific populations of *Aedes aegypti* (L.)**

Nancy M. Endersby-Harshman[1*•], Thomas L. Schmidt[1*], Jessica Chung[1], Anthony van Rooyen[2], Andrew R. Weeks[1,2], Ary A. Hoffmann[1]

[1]Pest and Environmental Adaptation Research Group, School of BioSciences, Bio21 Institute, 30 Flemington Rd, Parkville, The University of Melbourne, Victoria 3010, Australia

[2]cesar Pty Ltd, 293 Royal Parade, Parkville Victoria 3052, Australia

*Equal first authors
