## Supplemental Figure S1 for "Heterogeneous genetic invasions of three insecticide resistance mutations in Indo-Pacific populations of *Aedes aegypti* (L.)"

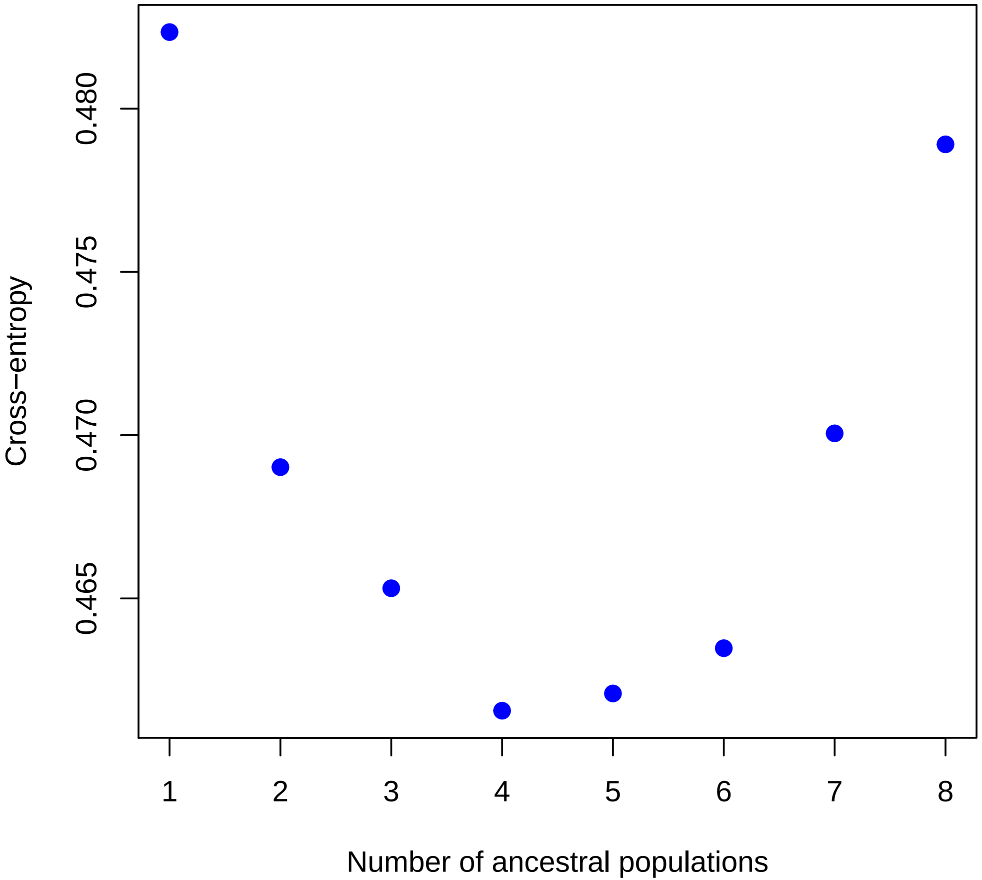

Cross−entropy
0.465
0.470
0.475
0.480
1
2
3
4
5
6
7
8
Number of ancestral populations

Supplementary Figure S1. Using the entire set of 50,569 SNPs, sparse nonnegative matrix factorization in LEA found *K* = 4 to be the optimal choice for *K*.

From:
Heterogeneous genetic invasions of three insecticide resistance mutations in Indo-Pacific populations of *Aedes aegypti (*L.)

Nancy M. Endersby-Harshman[1]*●, Thomas L. Schmidt[1]*, Jessica Chung[1], Anthony van Rooyen[2], Andrew R. Weeks[1,2], Ary A. Hoffmann[1]

[1]Pest and Environmental Adaptation Research Group, School of BioSciences, Bio21 Institute, 30 Flemington Rd, Parkville, The University of Melbourne, Victoria 3010, Australia
[2]cesar Pty Ltd, 293 Royal Parade, Parkville Victoria 3052, Australia

*Equal first authors
