## Supplemental Information for "Heterogeneous genetic invasions of three insecticide resistance mutations in Indo-Pacific populations of *Aedes aegypti* (L.)"

MOLECULAR ECOLOGY

*Equal first authors

### Table of Contents:


| | |
|---|---|
| Supplementary Table S1 | Page 2 |
| Supplementary Table S2 | Page 3 |
| Supplementary Information S1 | Page 7 |

MOLECULAR ECOLOGY

Table S1. Samples of *Aedes aegypti* collected from the Indo-Pacific region for studies of sodium channel (*Vssc*) mutations

| Collection date | Country | Collection site | Latitude | Longitude | n |
|---|---|---|---|---|---|
| Apr-2018 | Sri Lanka | Kiribathgoda | 6.980 | 79.925 | 24 |
| Apr-2016 | Indonesia | Bali – Denpasar Site 7 | -8.734 | 115.170 | 6 |
| Apr-2016 | Indonesia | Bali – Denpasar Site H | -8.717 | 115.176 | 12 |
| Apr-2016 | Indonesia | Bali – Denpasar Site 17 | -8.740 | 115.171 | 10 |
| Apr-2016 | Indonesia | Bali – Denpasar Site 11 | -8.739 | 115.168 | 9 |
| Apr-2016 | Indonesia | Bali – Denpasar Site 19 | -8.716 | 115.175 | 1 |
| Feb-2017 | Indonesia | Bali – Airport | -8.747 | 115.166 | 13 |
| Feb-2017 | Indonesia | Bali – North Kuta | -8.659 | 115.160 | 10 |
| Feb-2017 | Indonesia | Bali – Ubud | -8.507 | 115.261 | 6 |
| Apr-2015 | Singapore | Marsling Drive | 1.4416 | 103.773 | 3 |
| Apr-2015 | Singapore | Yishun Street | 1.423 | 103.834 | 4 |
| Apr-2015 | Singapore | Marsling Lane | 1.445 | 103.776 | 3 |
| Apr-2015 | Singapore | Farrer Road | 1.313 | 103.804 | 3 |
| Apr-2015 | Singapore | Jalan Chempadek | 1.371 | 103.830 | 4 |
| Apr-2015 | Singapore | Ang Mo Kio Ave | 1.372 | 103.844 | 1 |
| Apr-2015 | Singapore | Bukit Batok West Ave | 1.348 | 103.745 | 2 |
| Apr-2015 | Singapore | Clementi Ave | 1.320 | 103.768 | 4 |
| Apr-2015 | Singapore | West Coast Drive | 1.315 | 103.759 | 5 |
| May-2016 | Malaysia | Kuala Lumpur, Sunway FO | 3.070 | 101.605 | 16 |
| May-2016 | Malaysia | Kuala Lumpur, Hospital | 3.172 | 101.700 | 14 |
| May-2016 | Thailand | Bangkok – TBR | 13.717 | 100.739 | 10 |
| May-2016 | Thailand | Bangkok – WBK | 13.752 | 100.228 | 10 |
| May-2016 | Thailand | Bangkok – SKK | 13.679 | 100.722 | 7 |
| May-2016 | Thailand | Bangkok – JOY | 13.669 | 100.689 | 8 |
| May-2016 | Thailand | Bangkok – NAP | 13.723 | 100.753 | 5 |
| Aug-2015 | Vietnam | Tri Nguyen Island | 12.192 | 109.222 | 33 |
| Sep-2015 | Vietnam | Tri Nguyen Island | 12.192 | 109.222 | 28 |
| Sep-2015 | Vietnam | Nha Trang City | 12.243 | 109.189 | 36 |
| Feb-2016 | Taiwan | | 22.852 | 120.245 | 20 |
| Feb-2018 | New Caledonia | Quartier Latin | -22.276 | 166.445 | 6 |
| Feb-2018 | New Caledonia | Ctre Ville, Port Area | -22.272 | 166.439 | 12 |
| Feb-2018 | New Caledonia | Anse Vata | -22.300 | 166.445 | 4 |
| Feb-2018 | New Caledonia | Val Plaisance | -22.300 | 166.453 | 2 |
| Feb-2018 | Vanuatu | Joint Court | -17.742 | 168.320 | 10 |
| Feb-2018 | Vanuatu | Black Sands | -17.706 | 168.304 | 10 |
| Feb-2018 | Fiji (Nadi) | Nadi – Site 1 | -17.773 | 177.434 | 1 |
| Feb-2018 | Fiji (Nadi) | Nadi – Site 2 | -17.769 | 177.474 | 1 |
| Feb-2018 | Fiji (Nadi) | Nadi – Site 3 | -17.823 | 177.398 | 4 |
| Feb-2018 | Fiji (Nadi) | Nadi – Site 4 | -17.768 | 177.428 | 1 |
| Feb-2018 | Fiji (Nadi) | Nadi – Site 5 | -17.773 | 177.434 | 7 |
| Feb-2018 | Fiji (Nadi) | Nadi – Site 6 | -17.768 | 177.428 | 5 |
| Feb-2018 | Fiji (Nadi) | Nadi – Site 7 | -17.764 | 177.446 | 5 |
| Dec-2017 | Kiribati | | 1.984 | -157.369 | 20 |

MOLECULAR ECOLOGY

**Supplementary Table S2**

Details of the 80 *Ae. aegypti* used to test for genetic invasions.
These include some individuals previously sequenced by Schmidt et al. (2019), that have been aligned to the AaegL5 genome assembly (Matthews et al., 2018) used in this study.

| id | pop | location | geno | vssc | missingness | Profile GTC | Profile TGT |
|---|---|---|---|---|---|---|---|
| aeg01_Bali | Bali | Denpasar | GGTTCC | H1 H1 | 0.012127 | y | n |
| aeg02_Bali | Bali | Denpasar | GGTTCC | H1 H1 | 0.0152386 | y | n |
| aeg03_Bali | Bali | Denpasar | GGTTCC | H1 H1 | 0.0159566 | y | n |
| aeg04_Bali | Bali | Denpasar | GGTTCC | H1 H1 | 0.0172331 | y | n |
| aeg05_Bali | Bali | Denpasar | GGTTCC | H1 H1 | 0.0173129 | y | n |
| aeg06_Bali | Bali | Denpasar | GGTTCC | H1 H1 | 0.0193873 | y | n |
| aeg07_Bali | Bali | Denpasar | GGTTCC | H1 H1 | 0.019866 | y | n |
| aeg08_Bali | Bali | Denpasar | GGTTCC | H1 H1 | 0.0199457 | y | n |
| aeg09_Fiji | Fiji | Nadi | TTGGTT | H2 H2 | 0.0125259 | n | y |
| aeg10_Fiji | Fiji | Nadi | TTGGTT | H2 H2 | 0.0137227 | n | y |
| aeg11_Fiji | Fiji | Nadi | TTGGTT | H2 H2 | 0.013962 | n | y |
| aeg12_Fiji | Fiji | Nadi | TTGGTT | H2 H2 | 0.0144407 | n | y |
| aeg13_Fiji | Fiji | Nadi | TTGGTT | H2 H2 | 0.0145205 | n | y |
| aeg14_Fiji | Fiji | Nadi | TTGGTT | H2 H2 | 0.0255306 | n | y |
| aeg15_Fiji | Fiji | Nadi | TTTGTT | H2 H3 | 0.0209031 | n | y |
| aeg16_Fiji | Fiji | Nadi | TTTTTT | H3 H3 | 0.0289612 | n | n |
| aeg17_Kiribati | Kiribati | Kiribati airport | GGTTCC | H1 H1 | 0.0248923 | y | n |
| aeg18_Kiribati | Kiribati | Kiribati airport | TGTGTC | H1 H2 | 0.0142812 | y | y |
| aeg19_Kiribati | Kiribati | Kiribati airport | TGTGTC | H1 H2 | 0.0203447 | y | y |
| aeg20_Kiribati | Kiribati | Kiribati airport | TGTGTC | H1 H2 | 0.0250519 | y | y |
| aeg21_Kiribati | Kiribati | Kiribati airport | TTGGTT | H2 H2 | 0.0165949 | n | y |

| | | | | | | | |
|---|---|---|---|---|---|---|---|
| aeg22_Kiribati | Kiribati | Kiribati airport | TTGGTT | H2 H2 | 0.0201851 | n | y |
| aeg23_Kiribati | Kiribati | Kiribati airport | TTGGTT | H2 H2 | 0.0240147 | n | y |
| aeg24_Kiribati | Kiribati | Kiribati airport | TTGGTT | H2 H2 | 0.0245732 | n | y |
| aeg25_Malaysia | Malaysia | Kuala Lumpur | TGTGTC | H1 H2 | 0.0153183 | y | y |
| aeg26_Malaysia | Malaysia | Kuala Lumpur | TGTGTC | H1 H2 | 0.0155577 | y | y |
| aeg27_Malaysia | Malaysia | Kuala Lumpur | TGTGTC | H1 H2 | 0.019467 | y | y |
| aeg28_Malaysia | Malaysia | Kuala Lumpur | TGTGTC | H1 H2 | 0.0220999 | y | y |
| aeg29_Malaysia | Malaysia | Kuala Lumpur | TTGGTT | H2 H2 | 0.017632 | n | y |
| aeg30_Malaysia | Malaysia | Kuala Lumpur | TTGGTT | H2 H2 | 0.0183501 | n | y |
| aeg31_Malaysia | Malaysia | Kuala Lumpur | TTGGTT | H2 H2 | 0.032711 | n | y |
| aeg32_Malaysia | Malaysia | Kuala Lumpur | TTGGTT | H2 H2 | 0.0564864 | n | y |
| aeg33_NewCaledonia | New Caledonia | Noumea | TTGGTT | H2 H2 | 0.0173927 | n | y |
| aeg34_NewCaledonia | New Caledonia | Noumea | TTTGTT | H2 H3 | 0.0207436 | n | y |
| aeg35_NewCaledonia | New Caledonia | Noumea | TTTGTT | H2 H3 | 0.0248125 | n | y |
| aeg36_NewCaledonia | New Caledonia | Noumea | TTTGTT | H2 H3 | 0.0256901 | n | y |
| aeg37_NewCaledonia | New Caledonia | Noumea | TTTTTT | H3 H3 | 0.0157172 | n | n |
| aeg38_NewCaledonia | New Caledonia | Noumea | TTTTTT | H3 H3 | 0.0166746 | n | n |
| aeg39_NewCaledonia | New Caledonia | Noumea | TTTTTT | H3 H3 | 0.0254508 | n | n |
| aeg40_NewCaledonia | New Caledonia | Noumea | TTTTTT | H3 H3 | 0.026089 | n | n |
| aeg41_Singapore | Singapore | Singapore | GGTTCC | H1 H1 | 0.0497846 | y | n |
| aeg42_Singapore | Singapore | Singapore | GGTTCC | H1 H1 | 0.0775491 | y | n |
| aeg43_Singapore | Singapore | Singapore | TGTGTC | H1 H2 | 0.0358226 | y | y |
| aeg44_Singapore | Singapore | Singapore | TGTGTC | H1 H2 | 0.0366204 | y | y |
| aeg45_Singapore | Singapore | Singapore | TGTGTC | H1 H2 | 0.0368597 | y | y |
| aeg46_Singapore | Singapore | Singapore | TTGGTT | H2 H2 | 0.0280836 | n | y |
| aeg47_Singapore | Singapore | Singapore | TTGGTT | H2 H2 | 0.0292804 | n | y |
| aeg48_Singapore | Singapore | Singapore | TTGGTT | H2 H2 | 0.0361417 | n | y |

| | | | | | | | |
|---|---|---|---|---|---|---|---|
| aeg49_SriLanka | Sri Lanka | Colombo | TGTTTC | H1 H3 | 0.0258497 | y | n |
| aeg50_SriLanka | Sri Lanka | Colombo | TTGGTT | H2 H2 | 0.069491 | n | y |
| aeg51_SriLanka | Sri Lanka | Colombo | TTTGTT | H2 H3 | 0.011409 | n | y |
| aeg52_SriLanka | Sri Lanka | Colombo | TTTGTT | H2 H3 | 0.0148396 | n | y |
| aeg53_SriLanka | Sri Lanka | Colombo | TTTGTT | H2 H3 | 0.0243338 | n | y |
| aeg54_SriLanka | Sri Lanka | Colombo | TTTGTT | H2 H3 | 0.0251316 | n | y |
| aeg55_SriLanka | Sri Lanka | Colombo | TTTTTT | H3 H3 | 0.0203447 | n | n |
| aeg56_SriLanka | Sri Lanka | Colombo | TTTTTT | H3 H3 | 0.0400511 | n | n |
| aeg57_Taiwan | Taiwan | Kaohsiung City | GGTTCC | H1 H1 | 0.0224988 | y | n |
| aeg58_Taiwan | Taiwan | Kaohsiung City | TGTGTC | H1 H2 | 0.0147599 | y | y |
| aeg59_Taiwan | Taiwan | Kaohsiung City | TGTGTC | H1 H2 | 0.0183501 | y | y |
| aeg60_Taiwan | Taiwan | Kaohsiung City | GGTTTC | H1 H4 | 0.0142014 | y | n |
| aeg61_Taiwan | Taiwan | Kaohsiung City | TTGGTT | H2 H2 | 0.0210627 | n | y |
| aeg62_Taiwan | Taiwan | Kaohsiung City | TTGGTT | H2 H2 | 0.0493059 | n | y |
| aeg63_Taiwan | Taiwan | Kaohsiung City | TGTGTT | H2 H4 | 0.0109303 | y | y |
| aeg64_Taiwan | Taiwan | Kaohsiung City | TGTGTT | H2 H4 | 0.0143609 | y | y |
| aeg65_Thailand | Thailand | Bangkok | GGTTCC | H1 H1 | 0.013962 | y | n |
| aeg66_Thailand | Thailand | Bangkok | GGTTCC | H1 H1 | 0.0248923 | y | n |
| aeg67_Thailand | Thailand | Bangkok | TGTGTC | H1 H2 | 0.0214616 | y | y |
| aeg68_Thailand | Thailand | Bangkok | TGTGTC | H1 H2 | 0.0217808 | y | y |
| aeg69_Thailand | Thailand | Bangkok | TGTGTC | H1 H2 | 0.0303175 | y | y |
| aeg70_Thailand | Thailand | Bangkok | TTGGTT | H2 H2 | 0.0191479 | n | y |
| aeg71_Thailand | Thailand | Bangkok | TTGGTT | H2 H2 | 0.0197064 | n | y |
| aeg72_Thailand | Thailand | Bangkok | TTGGTT | H2 H2 | 0.0232966 | n | y |
| aeg73_Vanuatu | Vanuatu | Port Vila | GGTTCC | H1 H1 | 0.00989309 | y | n |
| aeg74_Vanuatu | Vanuatu | Port Vila | GGTTCC | H1 H1 | 0.0128451 | y | n |
| aeg75_Vanuatu | Vanuatu | Port Vila | GGTTCC | H1 H1 | 0.0130046 | y | n |

MOLECULAR ECOLOGY

| | | | | | | | |
|---|---|---|---|---|---|---|---|
| aeg76_Vanuatu | Vanuatu | Port Vila | GGTTCC | H1 H1 | 0.0141216 | y | n |
| aeg77_Vanuatu | Vanuatu | Port Vila | GGTTCC | H1 H1 | 0.0154779 | y | n |
| aeg78_Vanuatu | Vanuatu | Port Vila | GGTTCC | H1 H1 | 0.0159566 | y | n |
| aeg79_Vanuatu | Vanuatu | Port Vila | GGTTCC | H1 H1 | 0.0192277 | y | n |
| aeg80_Vanuatu | Vanuatu | Port Vila | GGTTCC | H1 H1 | 0.0197862 | y | n |

MOLECULAR ECOLOGY

***Supplementary Information S1: ddRAD library construction and filtering***

### ddRAD library construction

We applied the double-digest restriction-site associated DNA sequencing (ddRADseq: Peterson, Weber, Kay, Fisher, & Hoekstra (2012)) protocol for *Ae. aegypti* developed by Rašić, Filipović, Weeks, & Hoffmann (2014) to construct RAD libraries.

We performed initial digestions of 100 – 200 ng of genomic DNA in a 40 µL reaction, using 10 units each of MluCI and NlaIII restriction enzymes (New England Biolabs, Beverly MA, USA), NEB CutSmart buffer, and water. Digestions were run for 3 hours at 37°C with no heat kill step, and the products were cleaned with Ampure XP paramagnetic beads (Beckman Coulter, Brea, CA). These were ligated to modified Illumina P1 and P2 adapters overnight at 16°C with 1,000 units of T4 ligase (New England Biolabs, Beverly, MA, USA), followed by a 10-minute heat-deactivation step at 65°C. We performed size selection with a Pippin-Prep 2% gel cassette (Sage Sciences, Beverly, MA) to retain DNA fragments of 300/350 – 450 bp.

Final libraries were created by pooling eight 10 µL PCR reactions per library, each consisting of 1 µL size-selected DNA, 5 µL of Phusion High Fidelity 2× Master mix (New England Biolabs, Beverly MA, USA) and 2 µL of 10 µM standard Illumina P1 and P2 primers, run for 12 PCR cycles. These were cleaned and concentrated using an 0.8× concentration of Ampure XP paramagnetic beads (Beckman Coulter, Brea, CA) to make the final libraries. Between 40 and 65 individuals were included in each library, and each library was run on a single sequencing lane. Libraries were sequenced paired-end at either the Australian Genome Research Facility (AGRF, Melbourne, Australia) or the University of Melbourne (Pathology) on a HiSeq 2500 (Illumina, California, USA) in either High Output or Rapid Run modes, using 100 bp chemistry.

MOLECULAR ECOLOGY

### Filtering details

We used the process_radtags program in Stacks v2.0 (Catchen, Hohenlohe, Bassham, Amores, & Cresko, 2013) to demultiplex sequence reads and trim the reads to 80 bp in length. Using a 15-bp sliding window, low-quality reads were discarded if the average phred score dropped below 20. Single and paired reads were concatenated and aligned to the *Ae. aegypti* nuclear genome assembly AaegL5 (Matthews et al., 2018) with Bowtie2 (Langmead & Salzberg, 2012), using '--very-sensitive' alignment settings.

Initial filtering aimed to remove closely related mosquitoes and those with high levels of missing data. We used the Stacks ref_map pipeline to build individual Stacks catalogs for each of the 10 populations, from which we called genotypes at RAD stacks at a 0.05 significance level following Stacks v2.0 default parameters (Rochette, Rivera-Colón, & Catchen, 2019). We generated VCF files for each catalog with the Stacks program populations, and filtered it further using VCFtools (Danecek et al., 2011). SNPs were required to be biallelic, be present in 75% of mosquitoes in each population, have a minor allele count of > 2 (following Linck & Battey (2019)), and have less than 5% missing data.

We used SPAGeDi (Hardy & Vekemans, 2002) to calculate Loiselle's $k$ (Loiselle, Sork, Nason, & Graham, 1995) among mosquitoes within each population, identifying pairs with putative first-degree relatedness ($k \geq 0.1875$) and omitting related mosquitoes in order of missing data so that all remaining pairs had $k < 0.1875$.

This left 80 *Ae. aegypti* for downstream analyses. We generated a new Stacks catalog containing the 80 genotypes, and produced a VCF file which we filtered using the Stacks program populations and VCFtools. SNPs were required to be biallelic, be present in 75% of mosquitoes in each population, have a minor allele count of > 2, and have less than 5% missing data.
